# Molecular basis underlying male sterility in bHLH142 overexpressing rice

**DOI:** 10.1101/116996

**Authors:** Swee-Suak Ko, Min-Jeng Li, Yi-Jyun Lin, Hong-Xian Hsing, Ting-Ting Yang, Tien-Kuan Chen, Chung-Min Jhong, Maurice Sun-Ben Ku

## Abstract

Development of stable male sterility lines is essential for efficient hybrid seed production. We previously showed that knockout of *bHLH142* in rice (*Oryza sativa*) causes pollen sterility by interrupting tapetal programmed cell death (PCD). In this study, we demonstrated that overexpression of *bHLH142* (OE142) under the control of ubiquitin promoter also leads to male sterility in rice by triggering the premature onset of PCD. Protein of bHLH142 was found to accumulate specifically in the OE142 anthers. Overexpression of bHLH142 induced early expression of several key regulatory transcription factors in pollen development. In particular, the upregulation of EAT1 at the early stage of pollen development promoted premature PCD in the OE142 anthers, while its downregulation at the late stage impaired pollen development by suppressing genes involved in pollen wall biosynthesis, ROS scavenging and PCD. Collectively, these events led to male sterility in OE142. Analyses of related mutants further revealed the hierarchy of these pollen development regulatory genes. Thus, the findings of this study create a new method to generate genic male sterility in rice. Exploitation of this novel functionality of *bHLH142* would confer a big advantage to hybrid seed production.

**Highlight:** Overexpression of *bHLH142* leads to male sterility in transgenic rice due to early onset of tapetal PCD. This study creates a new method to generate male sterility in rice.

## Introduction

Rice (*Oryza sativa*) is one of the most important staple crops in the world, feeding almost half of the world’s population. Increase in rice production is urgently needed to keep pace with increasing population, especially in the face of drastic global climate change. Hybrid rice is considered the most promising strategy to increase grain yield increasing rice yields by 15–20% (Khush, 2013). By adopting hybrid technology, many countries have successfully increased per capita rice production (Zhang, 2011).

Heterosis in the F1 plants not only increases grain yield but also produces superior phenotypes in comparison with the parents with vigor in growth, good agronomic traits and pest resistance, etc. (Liu *et al.*, 2015). As rice is a self-pollinated crop, adoption of a stable male sterility female parent is critical to ensure the purity of F1 seeds. At present, two distinct hybrid systems, a three-line system and a two-line system, have been widely adopted in rice hybrid seed production. In the three-line system, cytoplasmic male sterility (CMS, A line) is the stable line. However, to maintain CMS seed stock, it is necessary to cross with a maintainer line (B line), which increases the production cost.

In the two line system, genic male sterility using photoperiod- or temperature-inducible male sterility mutant has the advantages of easy maintenance of seed stock and lower production cost, but it has the disadvantage of producing hybrid seeds of less uniformity due to environment fluctuation. Therefore, a better understanding of the mechanism underlying pollen development is important for developing new genic male sterility lines.

Rice anthers comprise four lobes and each lobe contains four layers of anther walls. The tapetum layer is in the innermost layer, providing nutrients and sporopollenin precursors for pollen development. Tapetal programmed cell death (PCD) at the right time is important for normal pollen development. In the anther, PCD is first detectable at meiosis (stage 8, S8), strong PCD signals occur at the young microspore stage (S9) (Li et al., 2006; Zhang and Wilson, 2009), and reduced PCD signals occur at the vacuolated pollen stage (S10) (Hu *et al.*, 2011). Functioning as polar secretory cells, the tapetum undergoes cellular degradation. Tapetal PCD subsequently triggers cytoplasmic shrinkage, breakdown of the nuclear membrane, oligonucleosomal cleavage of DNA, vacuole rupture, and swelling of the endoplasmic reticulum for release of mature pollen grains (Papini et al., 1999). Thus, tapetal PCD is an essential process for pollen maturation.

Pollen development involves a complex regulatory network. So far, several basic helix-loop-helix (bHLH) transcription factors (TFs) have been identified to play important roles in regulating tapetal PCD and pollen development. The roles of *UDT1* (*bHLH164*) (Jung *et al.*, 2005), *bHLH142 (TIP2)* (Fu *et al.*, 2014;Ko *et al.*, 2014), *TDR1 (bHLH5)* (Li *et al.*, 2006), and *EAT1 (DTD1, bHLH141)* (Ji *et al.*, 2013;Niu *et al.*, 2013) in rice pollen development have been characterized in the last decade. Similarly, *DYT1*, the homolog gene of *UDT1* in *Arabidopsis* (Zhang *et al.*, 2006) and *AMS* (Sorensen *et al.*, 2003), the homolog of *TDR1* in *Arabidopsis*, are functionally conserved in both dicots and monocots. In addition, the *UDT1* homolog in tomato, *ms10^35^* (*Solyc02g079810*), was also recently reported to be involved in pollen development (Jeong *et al.*, 2014). Another TF, GAMYB, is also known to play an important role in anther and aleurone layer development (Kaneko *et al.*, 2004;Tsuji *et al.*, 2006). According to current understanding of the pollen development regulatory network, UDT1 and GAMYB work in parallel to regulate pollen development and TDR1 acts downstream of UDT1 and GAMYB (Liu *et al.*, 2010). Our previous study showed that bHLH142 is located downstream of UDT1 and GAMYB and coordinates with TDR1 through protein-protein interaction to modulate *EAT1* transcriptional activity.

In addition, EAT1 interacts with TDR1 at a similar binding site to bHLH142 (Ko *et al.*, 2014). So far, the biological role of TDR1 in interacting with EAT1 remains unknown (Ji *et al.*, 2013;Ko *et al.*, 2014;Niu *et al.*, 2013). EAT1 directly regulates tapetal PCD via two aspartic proteases (AP37 & AP25) that activate cell death (Niu *et al.*, 2013).

AtTDF1 encodes a R2R3 MYB TF, which functions in callose dissolution (Zhu *et al.*, 2008). Similarly, the rice ortholog, OsTDF1 (MYB35), acts downstream of UDT1 and upstream of TDR, EAT1, OsMYB103 and Persistent Tapetal Cell 1 (PTC1) and it is essential for tapetal PCD (Cai *et al.*, 2015). PTC1 encodes a PHD-finger TF and controls tapetal PCD and pollen development and acts downstream of GAMYB (Li *et al.*, 2011) and TIP2 (bHLH142) (Fu *et al.*, 2014).

During anther development, ROS acts as a signal to promote tapetal PCD (Hu *et al.*, 2011;Yi *et al.*, 2016). The cellular ROS level is determined by the interplay between ROS-producing and ROS-scavenging mechanisms (Gapper and Dolan, 2006;Miller *et al.*, 2008). MADS3, a floral homeostatic C-class gene required for stamen identity, also regulates ROS scavenging during rice anther development. *MADS3* modulates ROS levels through positive transcriptional regulation of the promoter of metallothionein gene *MT-1-4b* (Hu *et al.*, 2011). The anthers of *mads3* mutant showed a strong ROS signal and a defect in pollen fertility (Hu *et al.*, 2011). On the other hand, defective Tapetum Cell Death 1 (DTC1) encodes a protein that contains a development and cell death (DCD) domain and KELCH repeats and acts as a key regulator of tapetum PCD by inhibiting ROS-scavenging activity through its interaction with metallothionein protein MT2b (Yi et al., 2016). Both MT-1-4b and MT2b act as ROS scavengers. Decreased expression of *MT2b* or *MT-1-4b* reduces scavenging activity and causes the accumulation of ROS molecules in rice roots (Steffens and Sauter, 2009) and anthers (Hu *et al.*, 2011). Therefore, a timely buildup of ROS to trigger tapetal PCD during pollen development is vital.

The pollen wall is composed of three layers: pollen coat, outer exine layer, and inner intine layer (Zhang *et al.*, 2016). Biosynthesis, secretion, and translocation of sporopollenin precursors are essential for pollen wall development. Synthesis of sporopollenin precursors is conducted in the tapetum, and *ACOS5, CYP703A, CYP704B, MS2*, etc., play major roles in this process. Ubish bodies transport tapetum-derived sporopollenin precursors to developing exine. Lipidic pollen exine is made of sporopollenin that is derived from the polymerization of fatty acid metabolites and phenolic acid (Ariizumi and Toriyama, 2011). In addition, MYB80/MYB103 is required for anther development in both *Arabidopsis* and rice (Higginson *et al.*, 2003;Zhang *et al.*, 2007). Male Sterility1 (MS1), a homeodomain (PHD) finger motif TF, regulates biosynthesis and secretion of pollen wall components in *Arabidopsis* (Wilson *et al.*, 2001;Yang *et al.*, 2007). A subsequent study by Li et al. (2011)found that *PTC1*, a *MS1* homolog in rice, is also essential for tapetal PCD and pollen development in rice (Li *et al.*, 2011). Several genes associated with rice pollen wall development have been identified by microarray analysis; these include *Cys protease* (*CP1;* Lee et al., 2004), a fatty acyl-CoA reductase homologous to *Arabidopsis MS2* (Aarts et al., 1997), lipid transfer proteins such as *C4* (Tsuchiya *et al.*, 1994), *C6* (Zhang *et al.*, 2010), YY1, BURP domain-containing proteins *(RA8* and *OsRAFTIN;* Jeon et al., 1999), and a P450 family member *CYP704B2* (Li et al., 2010). They were all down-regulated in the anther of the rice *ptc1* mutant (Li *et al.*, 2011). Moreover, mutagenesis studies suggest that *CYP703A2* (Yang *et al.*, 2014) and *CYP704B2* (Li *et al.*, 2010) are essential for pollen development, and their knockout lines exhibited impaired pollen development. *MS2* is essential for pollen wall biosynthesis by mediating the production of the conserved plastidial pathway for the production of fatty alcohols that are essential for pollen wall biosynthesis (Chen *et al.*, 2011;Shi *et al.*, 2011). Clearly, interruption of the functions of these genes resulted in abnormal pollen development.

In rice, *bHLH142* is specifically expressed in the anther and regulates tapetal PCD and pollen wall development, and knockout of *bHLH142* causes pollen sterility (Ko et al., 2014). To gain more insights into its functionality, in this study we generated transgenic lines overexpressing *bHLH142* under the control of maize ubiquitin promoter. We demonstrated that constitutive overexpression of *bHLH142* also leads to male sterility by triggering premature tapetal PCD. Our study provides a new method to generate genic male sterility in rice and possibly in other cereal crops too that may be suitable for agricultural application.

## Materials and Methods

### Constructs

The *bHLH142* (Os01g0293100) full length cDNA was PCR amplified using primers S80qPCR-F3_BamHI and S80FLcds-R2_BamHI (see online **Supplementary Table 1**) and the product is a 1373 bp BamHI fragment. This fragment was then digested with BamHI and ligated into pCAMBIA1390 backbone containing the maize ubiquitin promoter. Expression of the selection marker *HptII* gene that encodes hygromycin phosphotransferase was driven by cauliflower mosaic virus (CaMV) 35S promoter **(Supplementary Fig. S1E**). Another vector harboring fused bHLH142 and eGFP (Ubi::bHLH142- eGFP) was constructed to detect the tissue specificity of bHLH142 protein using eGFP. All constructs were confirmed by sequencing. The plasmids were separately transformed and selected by antibiotic. *Agrobaterium tumefaciens* strain EHA105 was used for transfection to calli of TNG67 background following the method described previously (Chan *et al.*, 1993).

### Plant material and growth conditions

Transformation of Japonica rice cultivar TNG67 was described previously (Ko et al., 2014). Primary transgenic lines were transplanted into soil and cultivated in the Academia Sinica-BCST greenhouse for genetically modified organisms, in Tainan, Taiwan.

### Southern blot analysis

Genomic DNA was extracted from leaf tissues using the cetyltrimethylammonium bromide (CTAB) method (Lee *et al.*, 2015). The genomic DNA was used for PCR and Southern blot analysis. Twenty micrograms of DNA was digested with HindIII and electrophoretically fractionated on a 0.8% agarose gel. Southern hybridization and detection were carried out using a digoxigenin-labeled *HptII* probe following the manufacturer’s instructions (Roche, http://www.rocheapplied-science.com).

### Histochemical staining

Transverse paraffin sections of anther were sectioned, deparaffined, rehydrated, and stained for starch with 2% I_2_/KI solution. Sudan Black B (0.3%, w/v; Sigma, Lot#MKBQ9075V) prepared in 70% ethyl alcohol was used to stain lipids, as described previously (Oliveira, 2015).

### TUNEL assay

To investigate the breakdown of tapetal programmed cell death, TUNEL assay was performed using the DeadEnd Fluorometric TUNEL system (Promega) as described previously (Ko *et al.*, 2014). Anther development stages from MMC (S7) to vacuolated pollen (S10) were collected.

### ROS staining and activity assay

Anthers of Wt and OE142 line #96 at various developmental stages were collected. Superoxide anion was quantified using a water-soluble tetrazolium salt reagent WST-1: Na,2-[4-iodophenyl] -3-[4-nitrophenyl]-5- [2,4-disulfophenyl]- 2H-tetrazolium), as described previously (Yi *et al.*, 2016).

### RNA isolation and qrt-pcr analyses

Rice (*Oryza sativa*) spikelets at different developmental stages, sporogenous cell (SC, S6), microspore mother cell (MMC, S7), meiosis (Mei, S8), young microspore (YM, S9), vacuolated pollen (VP, S10), pollen mitotic (PM, S11), and mature pollen at one day before anthesis (MP, S12), were collected for total RNA isolation, using LiCl2 method (Wang and Vodkin, 1994). One microgram of RNA was used to synthesize the oligo(dT) primed first-strand cDNA using the M-MLV reverse transcriptase cDNA synthesis kit (Promega). One microliter of the reverse transcription products was used as a template in the qRT-PCR reactions following previous protocols (Ko *et al.*, 2014). *Ubiquitin-like 5* (*UBQ5*, Os01g0328400) was used as an internal control for normalization of expression levels.

### Protein gel blot analysis

Total protein was extracted from newly matured leaves with Culture Cell Lysis Reagent (CCLR) buffer (100 mM K_2_HPO_4_, 100 mM KH_2_PO_4_, pH 7.8, containing 1% Triton X-100, 10% glycerol, 1 mM EDTA, and 7 mM 2-mercaptoenthanol). Protein concentration was measured using the Bio-Rad Protein Assay Kit with bovine serum albumin as a standard.

For western blot analysis, 80 μg total protein from each sample was loaded and separated by SDS-PAGE with a 12% acryamide gel and transferred onto polyvinylidene fluoride (PVDF) membrane for antibody probing. Antibodies against the rice bHLH142 and EAT1 were produced against the synthetic peptide (CSPTPRSGGGRKRSR) and (CELKILVEQKRHGNN), respectively. The following primary antibodies were used: Anti-bHLH142, rabbit polyclonal antibody (Genscript) at 1:4000 dilution, Anti-EAT1 rabbit polyclonal antibody (Genscript) at 1:2000 dilution, and Anti-eGFP rabbit polyclonal antibody (Yao-Hong Biotechnology, Cat#YH-80005) at 1:10000 dilution. Anti-Actin mouse monoclonal antibody (Sigma, A0480) at 1:2500 dilution was used as equal loading control.

### GFP fluorescence microscopy

Spikelet of the Wt, Ubi::bHLH142-eGFP and Ubi::GFP transgenic lines at the S9 to S10 stages were used for GFP florescence observation. GFP signal was recorded using a Zeiss LSM710 confocal microscope equipped with a T-PMT under an FITC filter at excitation of 488 nm and emission wavelength of 500–560 nm.

### RNA in-situ hybridization

Anthers of the non-transgenic Wt and OE142 at various developmental stages were collected and prepared in 10 μm thickness paraffin sections. Dig-labeled RNA probes of *bHLH142* and *EAT1* were cloned and prepared in advance. Hybridization protocols were as previously described (Ko *et al.*, 2014).

## Results

### bHLH142 overexpressing transgenic lines exhibit male sterility

For functional genomics studies, we generated transgenic rice lines overexpressing *bHLH142* under the control of the maize constitutive ubiquitin promoter in the japonica cultivar TNG67 (wild type, Wt) (**Fig. 1A**). More than 15 primary transgenic lines overexpressing *bHLH142* (OE142) were obtained and none of them produced viable seeds at the maturation stage. Genomic PCR confirmed T-DNA insertion in the OE142 lines (**Supplementary Fig. S1F**). With the exception of male sterility, the transgenic plants displayed Wt-like agronomic traits (**Fig.1**). All OE142 transgenic lines produced smaller anthers compared to the Wt (**Fig. 1C** and **Supplementary Fig. S1B**). Wt anthers dehisced normally during anthesis but OE142 anthers did not **(Fig. 1D right panel)**.

**Fig. 1.**
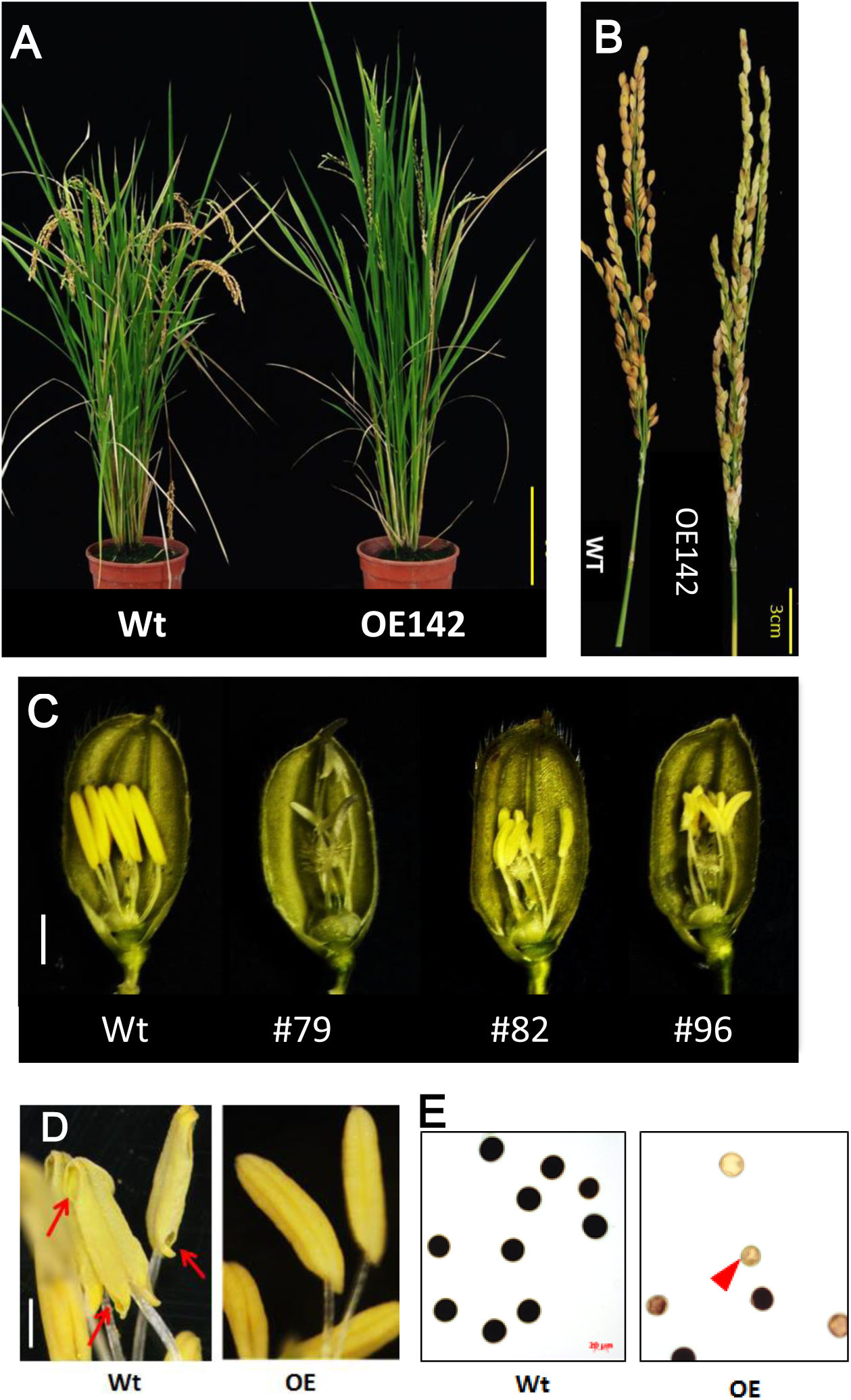
Overexpression of *bHLH142* (OE142) caused male sterility in rice. (A) Plant phenotype of wild-type (TNG67, Wt) and OE142 line #96 at seed maturation stage. (B) Panicles of Wt and OE142 at seed maturation stage. (C) Spikelets of Wt (in left) and several OE142 T0 lines at one day before anthesis. (D) Anthers dehiscence in the Wt but not in OE142 transgenic line. (E) Staining of pollen grains by 2% I2/KI solution in the Wt and OE142 line #96. Arrows show the dehisce anther (D). Arrowhead show infertile pollens (E). Scale bars: 20 cm (A), 3 cm (B), 2 mm (C), 20 μm (D, E).

Also, Wt pollen stained strongly by I_2_/KI but weaker staining was observed in the pollen of OE142 lines, indicative of low starch content **(Fig. 1E** and **Supplementary Fig. S2A)**. Finally, OE142 plants failed to produce viable seeds (**Fig. 1B** and **Supplementary Fig. S1D**).

To elucidate the defect in pollen maturation in OE142, detailed histological assays were carried out. OE142 anther produced less viable pollen grains, as demonstrated by I_2_/KI staining (**Fig. 1E** and **Supplementary Fig. S2A**). Moreover, OE142 anther showed a very weak Sudan Black staining of lipids compared to the Wt (**Supplementary Fig. S2B**). Histochemical staining analysis suggests that defect in starch and lipid synthesis in the OE142 anther may be caused by overexpression of *bHLH142.* Transverse section examination showed that OE142 anther entered the meiosis stage and the microspores were released into anther locules (**Supplementary Fig. S3**). However, abnormal anther development in OE142 was observed at the vacuolated pollen stage (S10) where epidermal layer was not thickened. Degeneration of OE142 pollen was observed at the pollen mitotic stage (S11) (**Supplementary Fig. S3**). At the anther maturation stage, Wt showed thickening of endothecial cell layers, ready for anther dehiscence (**Supplementary Fig. S2C**) but OE142 endothecial cell layers remained thin and no anther dehiscence took place (**Supplementary Fig. S2C, S3**). Finally, severely degenerated pollen grains were observed in OE142 at the anther maturation stage (**Supplementary Fig. S2C, S3**).

### OE142 has premature onset of tapetal PCD

As defect in pollen development was observed in OE142 anther (**Fig. 1**, and **Supplementary Fig. S1-S3**), we suspected that overexpression of *bHLH142* might have altered tapetal PCD, which is responsible for tapetum degeneration during maturation (Papini *et al.*, 1999). Therefore, TUNEL assay was performed to detect DNA fragmentation in the anthers of OE142 line in comparison to Wt. As shown in **Fig. 2**, Wt exhibited a normal tapetal PCD signal starting from meiosis-II stage (S8b), which was increased at the young microspore stage (S9). However, premature onset of tapetal PCD was clearly observed in the OE142 anthers, which started at stage S8a with the highest DNA fragmentation signal occurring at S8b, but reduced PCD at S9 (**Fig. 2**). The corresponding TUNEL differential image contrast (DIC) images showing the anatomy of anther are presented in **Supplementary Fig. S4**. These data indicate that overexpression of *bHLH142* triggered premature onset of tapetal PCD at S8a. However, OE142 lost timely tapetal PCD at S9 that is critical for releasing nutrients to nurture microspore development, leading to defected pollen maturation.

**Fig. 2.**
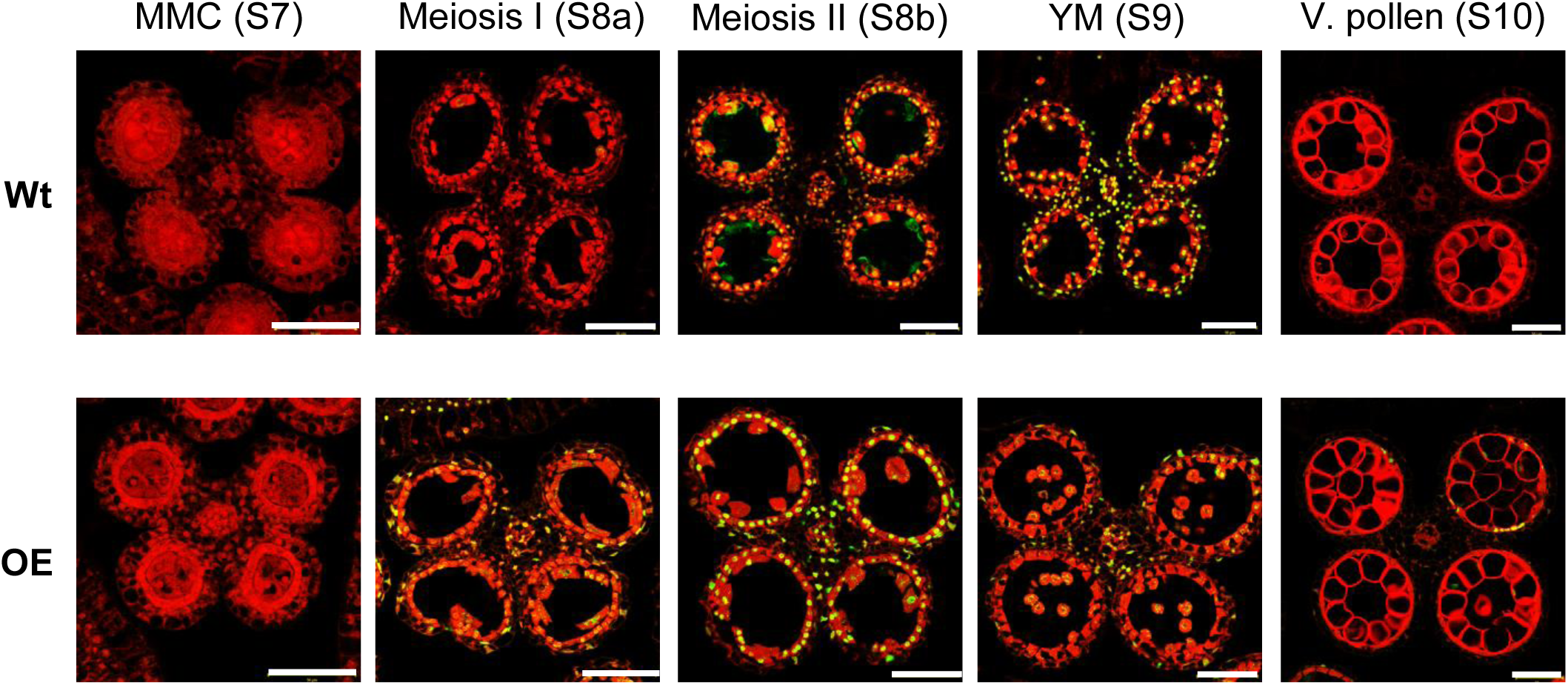
TUNEL assay showing premature onset of tapetal programmed cell death in the OE142 anther. DNA fragmentation signals (yellow fluorescence) started at the meiosis II stage (S8b) and exhibited obvious positive TUNEL signals at the young microspore stage (S9) in the Wt (upper panel). Early DNA fragmentation signal was observed in the tapetum of OE142 at meiosis I (S8a), and increased TUNEL positive signals occurred at the meiosis II (S8b) stage (lower panel). The red signal exhibits propidium iodide staining, and the yellow fluorescence is the merged signal from TUNEL (green) and propidium iodide staining (red). Scale bars: 50 μm.

### Molecular changes in OE142

Three OE142 lines with varying expression levels (#79, #82 and #96) were propagated vegetatively for further molecular studies (Fig. 1C). Southern blot hybridization analysis with *HptII* probe indicated that T-DNA insert was integrated into the rice genome at 3, 1, and 3 copies in the OE142 lines #79, #82, and #96, respectively (Fig. 3A). Real time PCR analysis further showed that the *bHLH142* transcript was 3.3, 9.5 and 51.7 fold higher in the anthers of these respective lines, compared to that of Wt anther (Fig. 3B). The OE142 line #96 expressed most abundant *bHLH142* mRNA and was therefore used for further molecular characterization, unless otherwise indicated. Irrespective of insertion copy number or *bHLH142* transcript abundance, all three OE lines failed to produce fertile pollen grains as a result of defect in pollen viability. This result implies that a proper expression level of *bHLH142* at the right stage is critical for maintaining normal pollen development in rice.

**Fig. 3.**
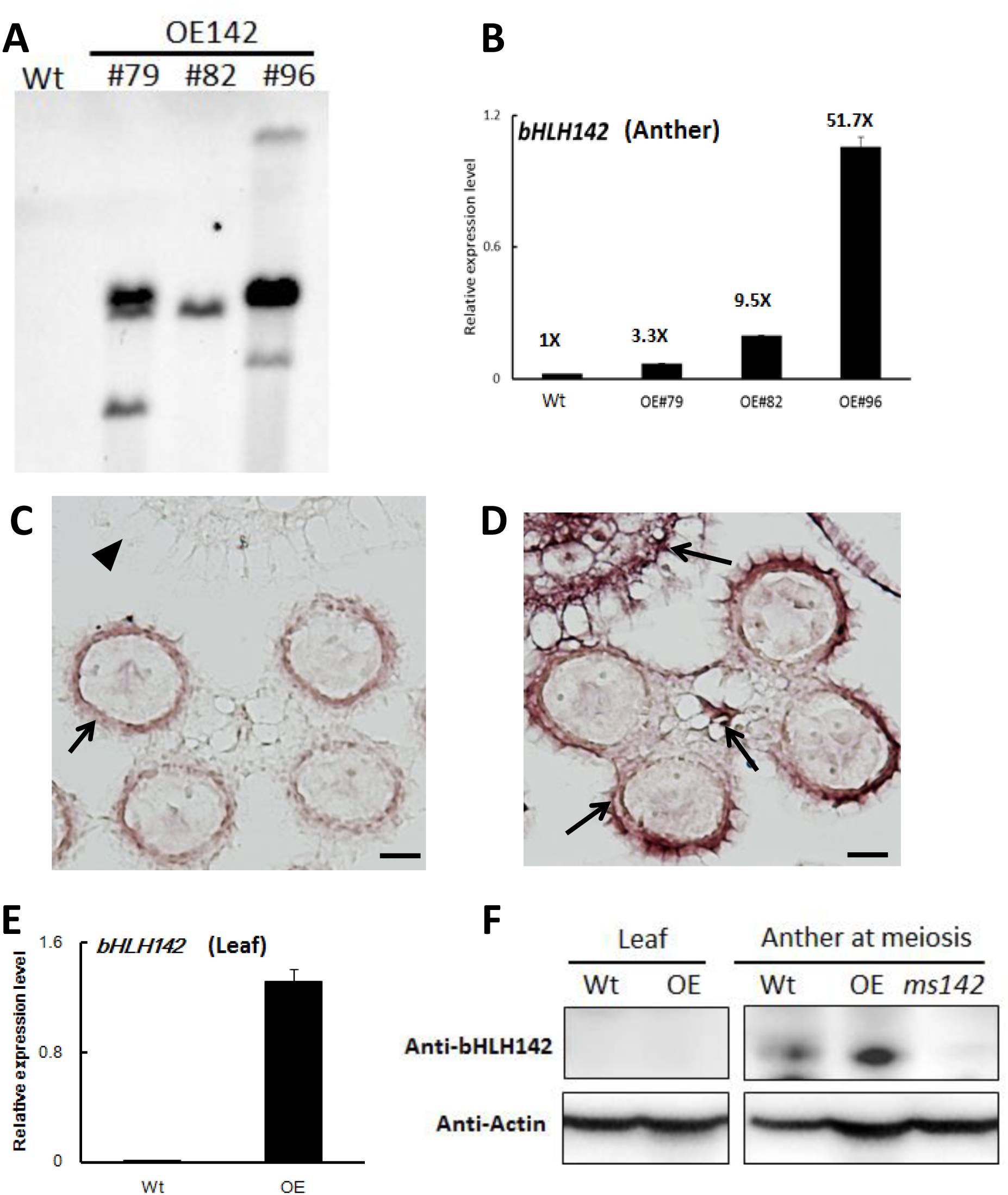
Expression patterns of bHLH142 mRNA and protein in the OE142 transgenic line. (A) Southern blot indicated T-DNA insertion number in the OE142 transgenic lines. Twenty micrograms of DNA from each line was digested with *HindIII* and fractionated on 0.8% agarose gel. Southern blot was carried out using a digoxigenin-labeled *HptII* probe. (B) qRT-PCR showed up-regulation of *bHLH142* transcript in the anther of OE lines at young microspore stage. (C) RNA ISH of *bHLH142* antisense probe hybridization in the Wt anther at meiosis stage (S8). (D) RNA ISH of *bHLH142* antisense probe hybridization in OE142 anther at meiosis stage (S8). (E) Overexpression of *bHLH142* significantly increased *bHLH142* transcripts in the leaves of OE142 as analyzed by qRT-PCR. (F) Protein of bHLH142 was not expressed in the leaves of OE142 as analyzed by western blot analysis. It was specifically expressed in the anthers. KO mutant, *ms142*, was included as a negative control for western blotting of bHLH142. Error bars indicate SD of mean from three replicates (B, E). Arrows indicated ISH positive signals in the anther walls (C); ISH positive signals in the hulls, vascular bundle, and anther walls of OE142 (E). The arrowhead indicates no ISH signal in the hull of Wt. Scale bars: 20 um (C, D).

### bHLH142 protein is specifically expressed in the anther

Previous RNA *in situ* hybridization (ISH) analysis indicated that *bHLH142* is tissue specifically expressed in the anthers of Wt at S7 to S9 but not in the leaf (Fu *et al.*, 2014;Ko *et al.*, 2014). RNA ISH data indicated that *bHLH142* transcript was localized specifically in the tapetum, middle layer, and meiocytes of the Wt (**Fig. 3C**). However, in OE142 transgenic line, *bHLH142* transcript was detected in leaf tissue (**Fig. 3E**) and anther (**Fig. 3B**). Moreover, RNA ISH analysis further indicated that *bHLH142* transcript is constitutively expressed in the hulls, anther walls, vascular bundle, and meiocytes of OE142 (**Fig. 3D**). Surprisingly, western blot analysis using anti-bHLH142 antibody showed that bHLH142 protein is only present in the anther but absent in the leaf of OE142 (**Fig. 3E**), suggesting post-transcriptional regulation of its expression. In addition, using Ubi::bHLH142-GFP transgenic plants generated in this study we demonstrated that GFP fluorescent signal was detected only in the anther but not in the hull of the transgenic line (**Fig. 4**). Consistently, western blot analysis of various tissues from Ubi::GFP and Ubi::bHLH142-GFP plants further demonstrated that GFP protein is only present in the anther but not in the leaf, hull or seed of Ubi::bHLH142-GFP transgenic line (**Fig. 4C**, right panel). Transgenic line overexpressing Ubi::GFP served as a good positive control showing constitutive expression of GFP protein in all tested organs (**Fig. 4C**, left panel). These results show that both Ubi::bHLH142 (OE142) and Ubi::bHLH142-GFP constructs drove the expression of bHLH142 protein specifically in the anther. Clearly, bHLH142 is expressed in an anther specific manner in OE142. Moreover, both Ubi::bHLH142 (OE142) and Ubi::bHLH142-GFP transgenic lines showed a similar male sterility phenotype, presumably due to the overexpression of bHLH142.

**Fig. 4.**
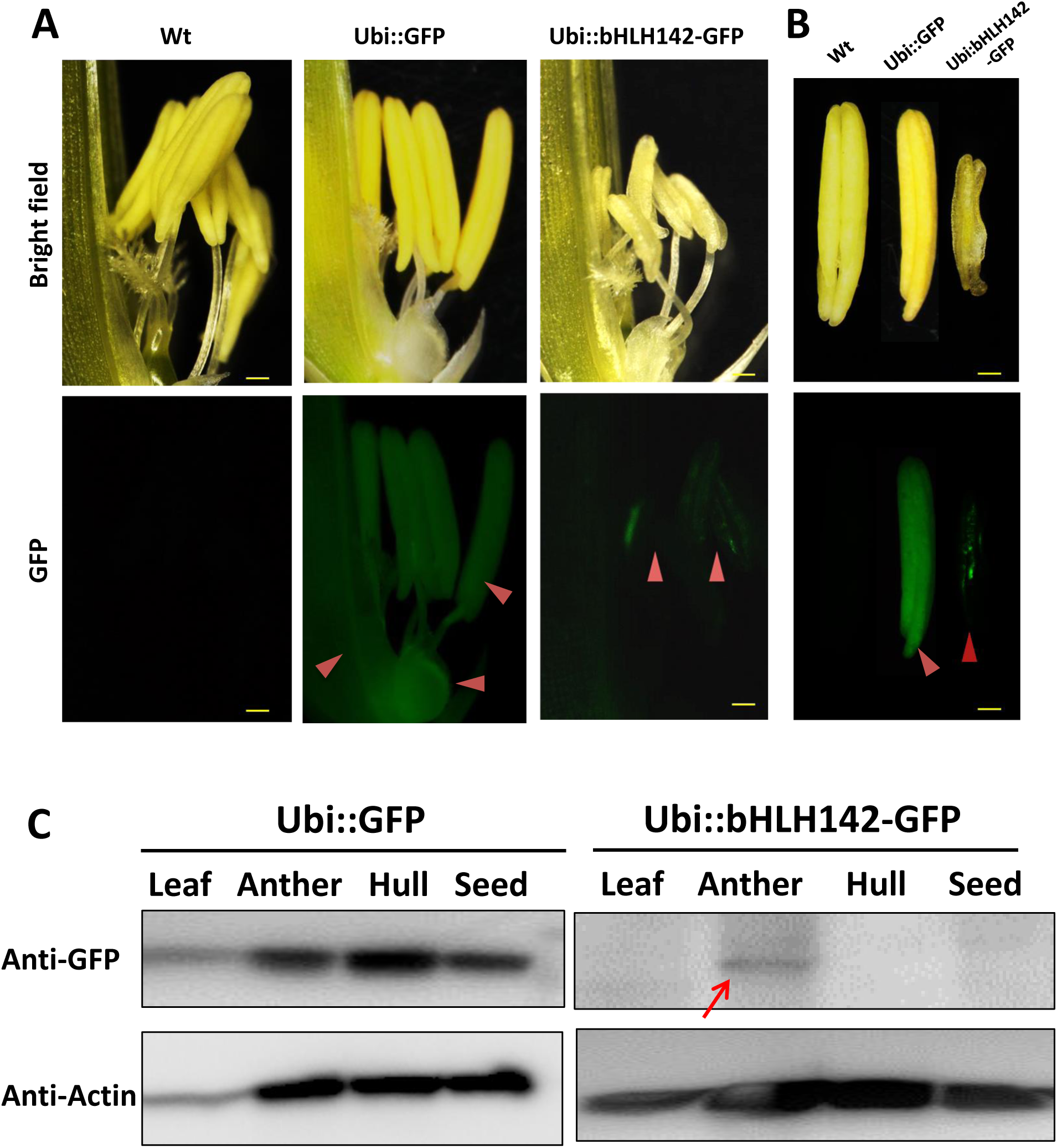
Protein of bHLH142 is specifically expressed in the anther. GFP signals observed under fluorescence microcopy for spikelet (A) and anthers (B). Upper panel shows bright field and lower panel shows GFP signals. Arrowheads show GFP signals in the spikelets and anthers of transgenic lines. Scale bars: 50 μm (A), 20 μm (B). (C) Western blotting using anti-eGFP antibody indicated bHLH142 protein is tissue specifically expressed in the anther of Ubi::bHLH142-GFP plant only. Transgenic rice overexpressing GFP as driven by ubiquitin promoter was included as a control. Anthers at vacuolated pollen stage were collected for protein isolation.

### Overexpression of bHLH142 alters transcriptional regulation of several known genes related to pollen development

To understand whether the pollen development regulatory network in OE142 was altered, qRT-PCR analysis of several of the known regulatory TFs that are involved in pollen development was carried out. As the expression of *bHLH142* in the OE142 lines was driven by the strong constitutive ubiquitin promoter, its mRNA expression in the OE142 lines was consistently upregulated throughout all stages of anther development (**Fig. 5A**). Interestingly, the expression of *GAMYB, UDT1 (bHLH164)*, and *MYB35 (TDF1)* was also upregulated in the OE142 anthers (**Fig. 5**). *TDR1 (bHLH5)* was upregulated at the early stage but then downregulated at the meiosis stage (S8) in OE142 (**Fig. 5E**). Similarly, *EAT1* was upregulated at the early meiosis stages from S6 to S8 but strongly suppressed after reaching the young microspore stage (S9) in the anthers of OE142 (**Fig. 5F**). In addition, the expression of *MYB80* was found downregulated at meiosis onwards (**Fig. 5G**). Similar to *MYB80, PTC1*, a key regulator of tapetal PCD and pollen wall biosynthesis (Li *et al.*, 2011), declined significantly to a negligible amount at S9 in the OE142 anthers (**Fig. 5H**). Clearly, constitutively overexpressing *bHLH142* alters the expression of the key regulatory TFs associated with pollen development.

**Fig. 5.**
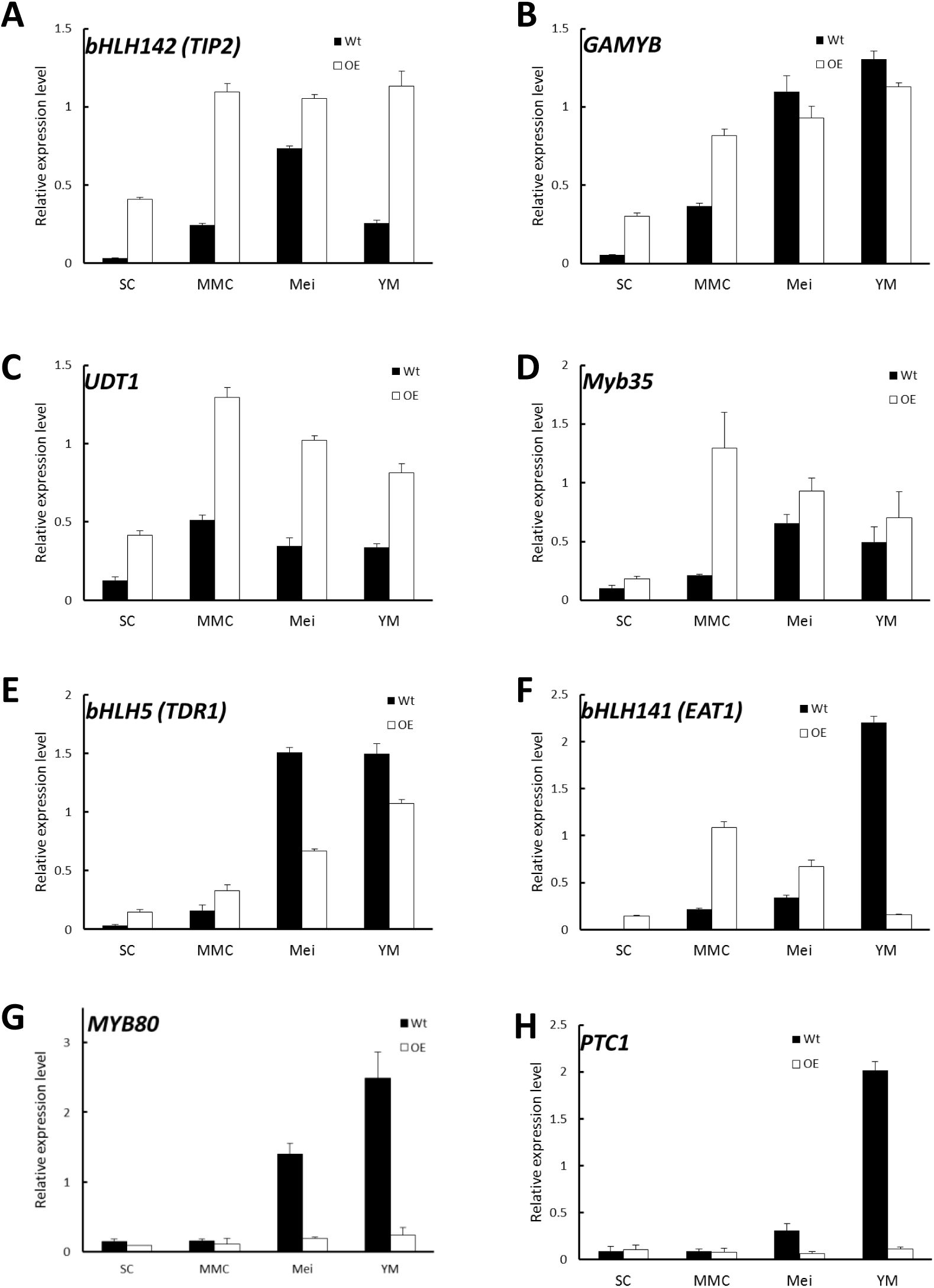
Overexpression of *bHLH142* altered expression in some transcription factors involved in pollen development. SC, sporogenous cell, S6; MMC, microspore mother cell, S7; Mei, meiosis, S8; YM, young microspore, S9. Error bars indicate the SD of mean from three replicates.

### Overexpression of bHLH142 downregulates PCD related functional genes

EAT1 is known to trigger tapetal PCD by regulating the expression of two *Aspartic Proteases (AP37, AP25)* at the young microspore stage (Niu *et al.*, 2013). In OE142 anthers, *EAT1* was significantly downregulated at S9 (**Fig. 5F**). TUNEL assay indicated premature onset of tapetal PCD in OE142 anthers (**Fig. 2**), which was correlated with the upregulation of *EAT1* before the meiosis stage (**Fig. 5F**). Slightly higher expression of *AP37* and *AP25* at the early stages of OE142 anther development was observed (**Fig. 6**). Normally, Wt rice exhibits the strongest expression of *EAT1, AP37, AP25*, and *CP1* at S9 to turn on timely tapetal PCD (**Fig. 5F, 6**). Thus, the negligible expression levels of these marker genes associated with PCD at S9 in the anthers of OE142 further supports the notion that decreased expression of these tapetal functional genes might disrupt timely tapetal PCD (**Fig. 2**). Consistently, reduced expression of these two proteases (*AP37* and *AP25)* coincided with the reduction of *EAT1* mRNA in OE142 at S9 (**Fig. 5F, 6**). Thus, collectively these results further supported the previous finding that EAT1 regulates *AP37* and *AP25* (Niu *et al.*, 2013).

**Fig. 6.**
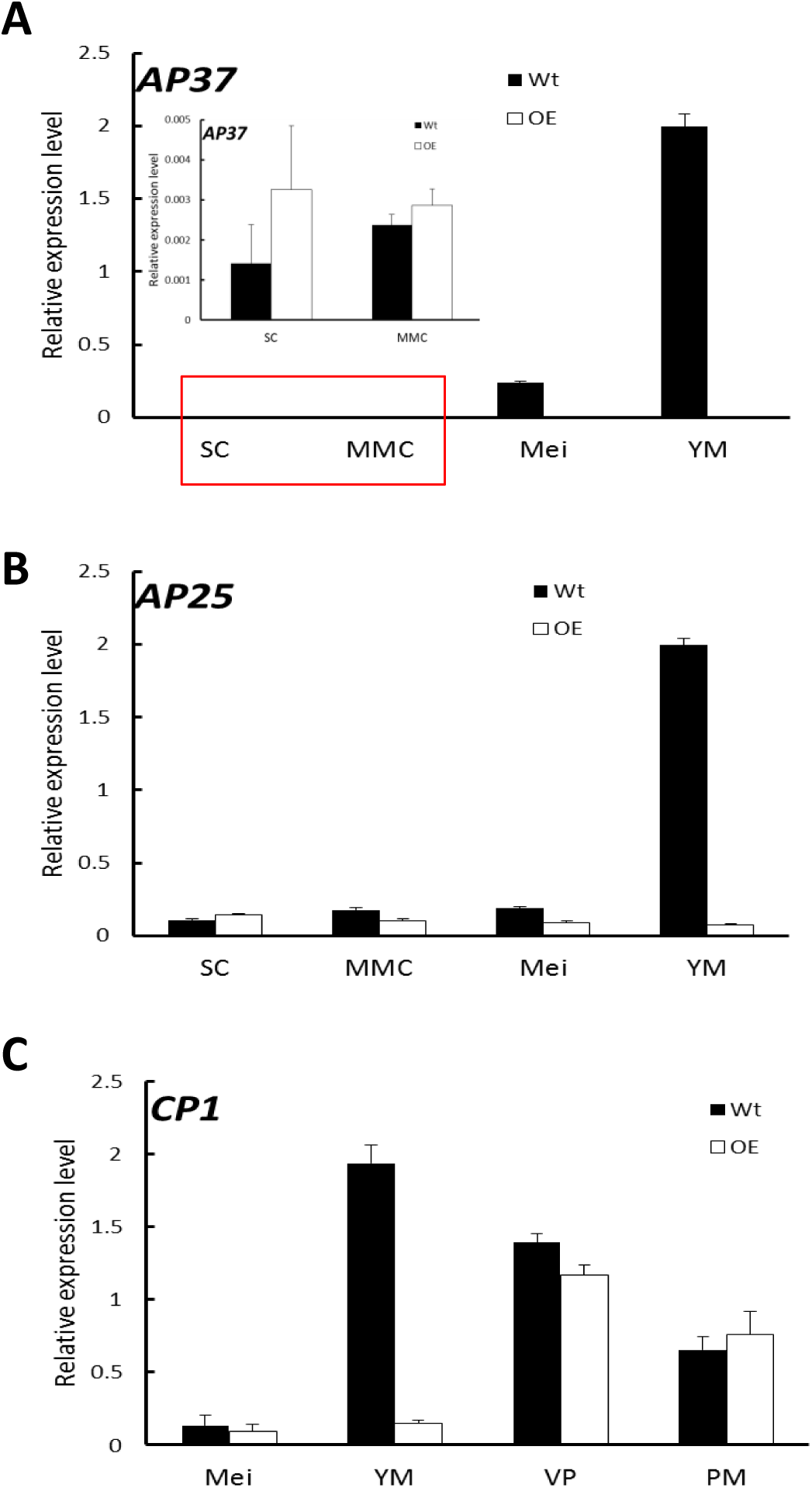
Overexpression of *bHLH142* altered expression patterns in tapetal PCD associated genes. (A) Gene expression of *AP37.* Red area is magnification. (B) Gene expression of *AP25.* (C) Gene expression of *CP1.* Abbreviations are as described in the legend for Fig. 5.

### OE142 has altered ROS metabolism in the anther

Timely accumulation of ROS is essential to induce PCD during tapetum degeneration (Hu *et al.*, 2011;Yi *et al.*, 2016). The premature onset of tapetal PCD as shown in **Fig. 2** prompted us to hypothesize that OE142 may have altered ROS metabolism in the anthers. Therefore, quantitative measurement of superoxide ion using WST-1 was performed in the anthers of the Wt and OE142 line at various developmental stages. Normally, ROS accumulates more at S8 to S9 to trigger tapetal PCD (Xie *et al.*, 2014). Our results showed that the Wt anthers accumulated the highest level of superoxide anions during the meiosis stage; however, OE142 had significantly lower level of superoxide anions compared to the Wt. In addition, the anthers of OE142 accumulated more superoxide anions at the later stage of anther development (**Fig. 7A**) that might be toxic for OE142 anther development. A previous study suggested that tapetal PCD requires timely and precise control of ROS levels (Xie *et al.*, 2014). We therefore compared the expression of rice ROS scavenging associate genes in OE142 at various stages of anther development. Our qRT-PCR analysis indicated that OE142 reduced the expression of *MADS3* and *MT2b* (**Fig. 7**). Earlier studies suggest that *MADS3* involved in ROS metabolism to trigger PCD (Hu *et al.*, 2011). Our qRT-PCR results indicated that *MADS3*, and *MT2b* were significantly downregulated in the anthers of OE142 (**Fig.7B, C**), which is consistent with the higher ROS accumulation in OE142 as compared to the Wt (**Fig. 7A**). Taken together, these results suggest that decreased ROS scavenging activity in OE142 anthers affects ROS metabolism and initiation of synchronized PCD, resulting in defective pollen grains (**Fig. 1E**). Based on the fact that *EAT1* expression is downregulated at the S9 stage in OE142 anthers, we contemplated whether *EAT1* may play an important regulatory role in ROS metabolism. To verify the possible gene hierarchy in this regulatory process, qRT-PCR analyses of the expression of ROS marker genes in *eat1* (Tos17 mutant) anthers were carried out. The results indicated that *MADS3 and MT2b were* downregulated in *eat1* mutant (**Fig. 7D, E**), implies that *MADS3* and *MT2b* genes might be located downstream of the *EAT1* regulatory network. Taken together, these data suggest that overexpression of *bHLH142* causes downregulation of *EAT1* at the late stage of anther development, which in turn alters the expression of ROS scavenging genes with decreased scavenging activity and accumulation of ROS molecules, leading to defected male gametophyte development.

**Fig. 7.**
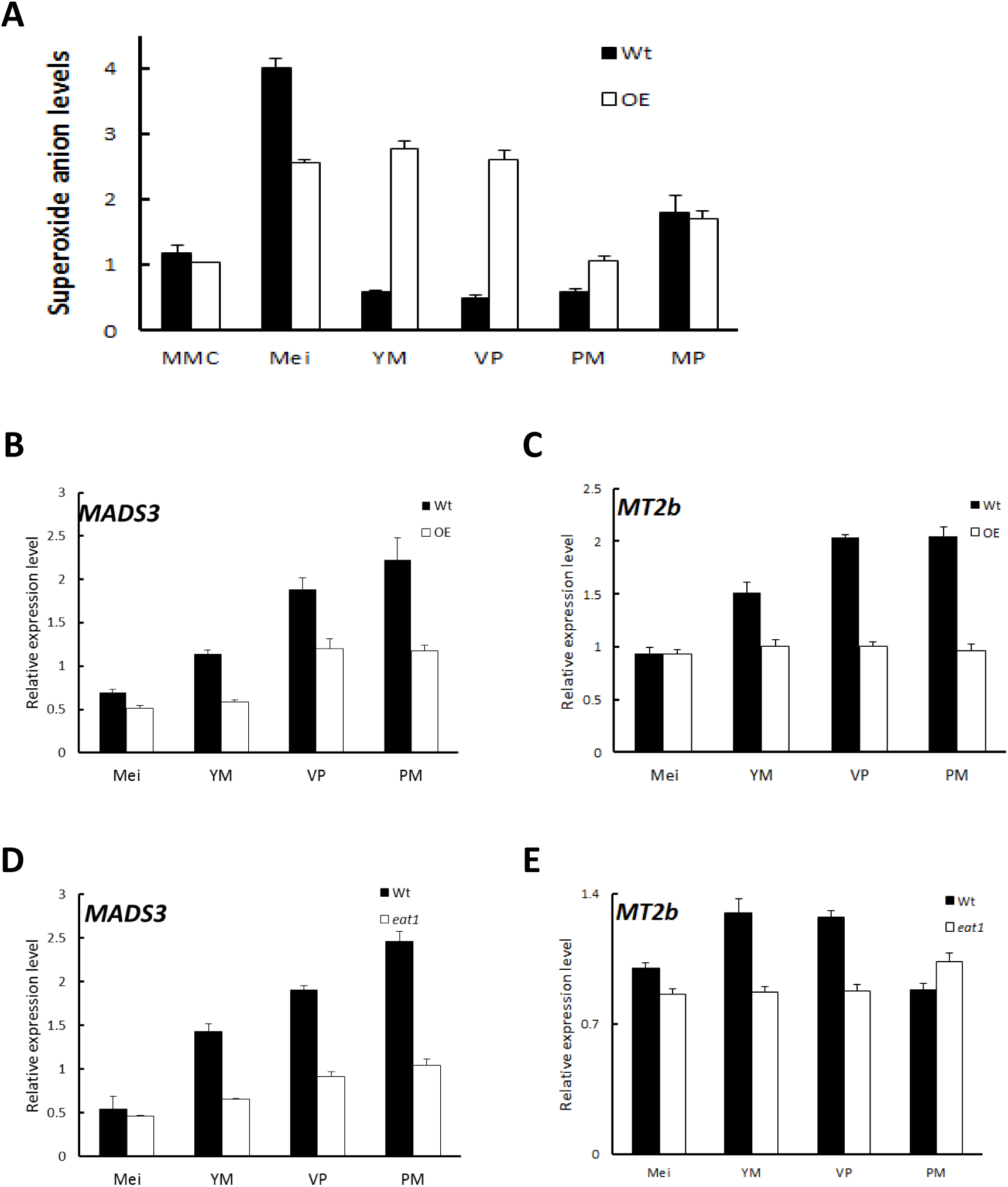
Overexpression of *bHLH142* altered superoxide anion accumulation and the expression of ROS-associated genes. (A) Alter superoxide anion levels in the anthers of OE142. (B) Comparison of Wt and OE142 using qRT-PCR to analyze gene expression patterns of *MADS3* and *MT2b* (C). (D-E) Mutagenesis analysis indicated gene hierarchy of *MADS3* and *MT2b* locate downstream of *EAT1.* Wild type for *eat1* is in Hitomebore background. Abbreviations are as described in the legend of Fig. 5.

### Overexpression of bLHL142 impairs sporopollenin biosynthesis

Lipidic exine synthesis is an important component of the pollen wall in rice and *Arabidopsis* (Yang *et al.*, 2007). The anthers of OE142 were weakly stained by the lipid specific dye Sudan Black compared to that of Wt (**Supplementary Fig. S2B**). Moreover, the TF *PTC1* was downregulated in OE142 (**Fig. 5H**). Several lipid transfer proteins were also downregulated in *ptc1* anthers (Li *et al.*, 2011). Therefore, the expression of these marker genes related to pollen sporopollenin biosynthesis was monitored by real time PCR during OE142 pollen development. The results demonstrated that overexpression of *bHLH142* sharply reduced the expression of these genes related to sporopollenin and pollen wall biosynthesis. The transcripts of *Cyp703A3, Cyp704B2, MS2*, and *C4* were almost not detectable in OE142. The expression of *C6* was also down-regulated in the anthers of OE142 at the late stage of development (**Fig. 8**). Our analyses with rice *tdr1* and *eat1* mutants also indicated that *MYB80* was downregulated at S9. Similarly, *PTC1*, a key regulator of sporopollenin biosynthesis, was significantly downregulated in the *eat1* anther (**Supplementary Fig. S5**). Taken together, these results support the idea that both *MYB80* and *PTC1* regulate sporopollenin biosynthesis in both monocots and dicots. Thus, overexpressing *bHLH142* caused downregulation of *EAT1* at S9, which might severely inhibit *MYB80* and *PTC1* and reduce sporopollenin gene expression (**Fig. 8**) and interrupt normal sporopollenin biosynthesis with defected pollen wall in OE142 transgenic lines.

**Fig. 8.**
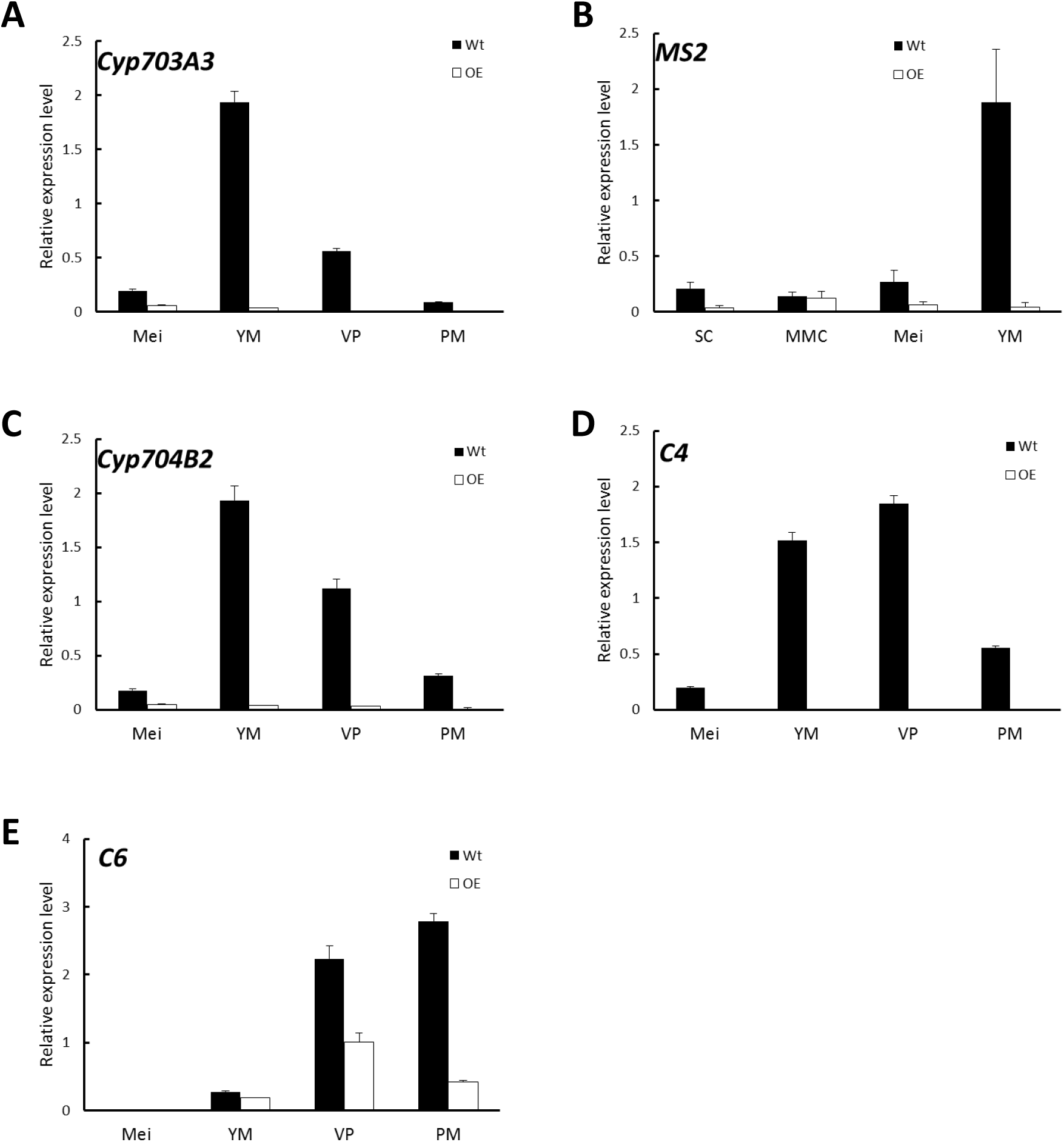
Overexpression of *bHLH142* altered the expression of genes associated with pollen wall biosynthesis. Comparison of Wt and OE142 using qRT-PCR to analyze gene expression patterns of *Cyp703A3* (A), *MS2* (B), *Cyp704B2* (C), *C4* (D), and *C6* (E). Abbreviations are as described in the legend of Fig. 5.

### OE142 anther exhibits parallel changes in EAT1 transcript and protein

In this study, we found that *EAT1* was upregulated at stages S6 to S8 but then downregulated at S9 in the anthers of OE142 (**Fig. 5F, 9A**). To understand the spatial and temporal expression patterns of *EAT1* in OE142, we carried out RNA ISH hybridized *EAT1* Dig-labeling probe in the anthers of Wt vs. OE142 at S8a and S9. The results revealed that *EAT1* mRNA was highly expressed in the tapetum, middle layer, meiocycte, microspore, vascular bundle, and hull of the Wt at S9 (**Fig. 9B**). However, ISH positive signal of *EAT1* was strong in the anthers of OE142 at early meiosis (S8a), but significantly reduced to a negligible level at YM (S9). The ISH results support our *EAT1* qRT-PCR data (**Fig. 5F**), providing a clear picture of the *in vivo* transcriptional map of *EAT1.*

**Fig. 9.**
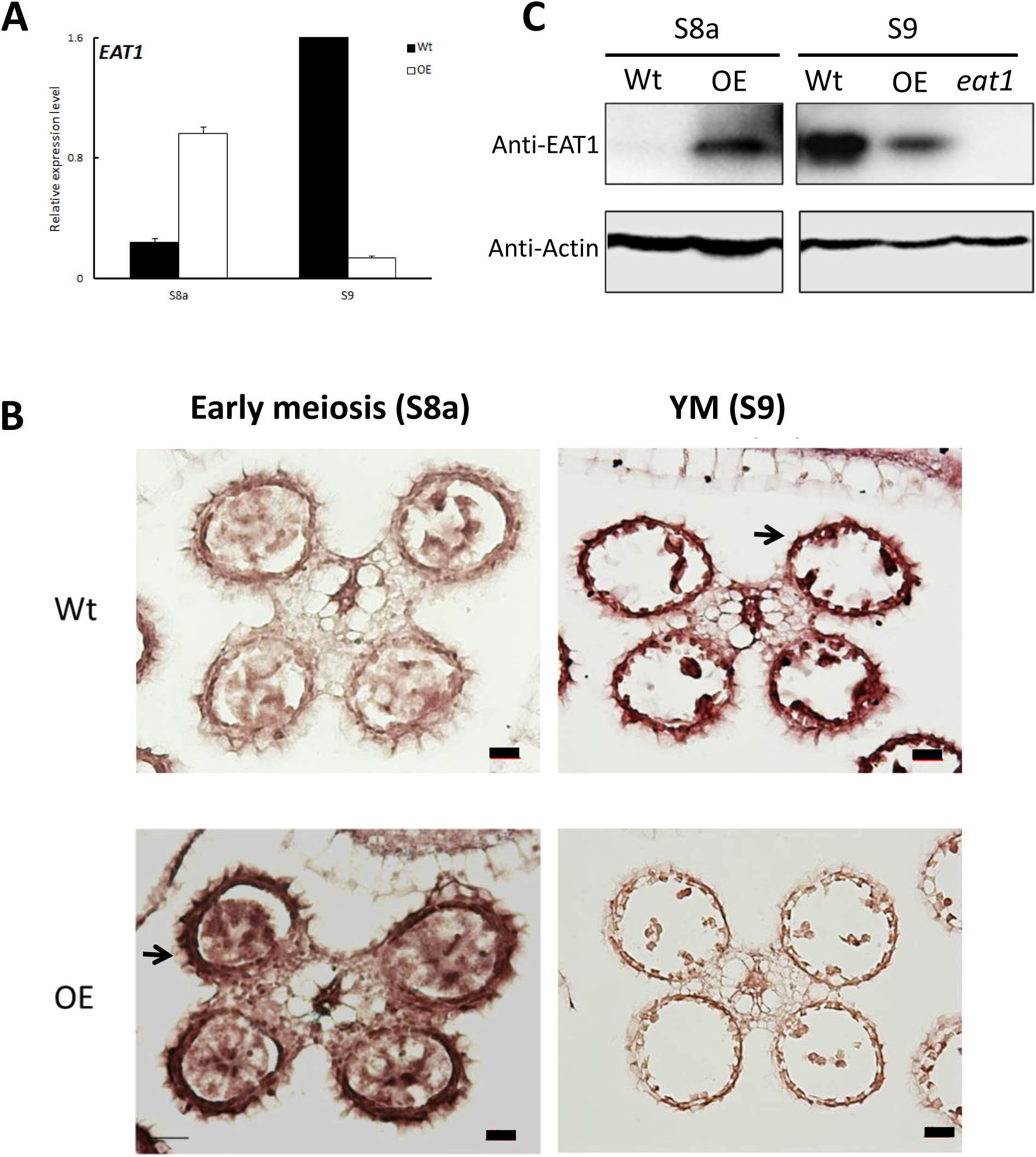
Transcript and protein levels of EAT1 were upregulated at the early stage and then downregulated at later stage of anther development in OE142. (A) qRT-PCR revealed upregulation of *EAT1* at (S7) and downregulation at YM (S9). (B) RNA ISH hybridization to EAT1-antisense probe in the Wt and OE142 anthers at stages S8 to S10. (C) Western blotting showed premature expression of EAT1 protein at MMC and downregulation at YM in OE142. The knockout mutant *eat1* was used as a negative control. Abbreviations are as described in the legend of Fig. 5. Scale bars: 20 μm.

**Fig. 10.**
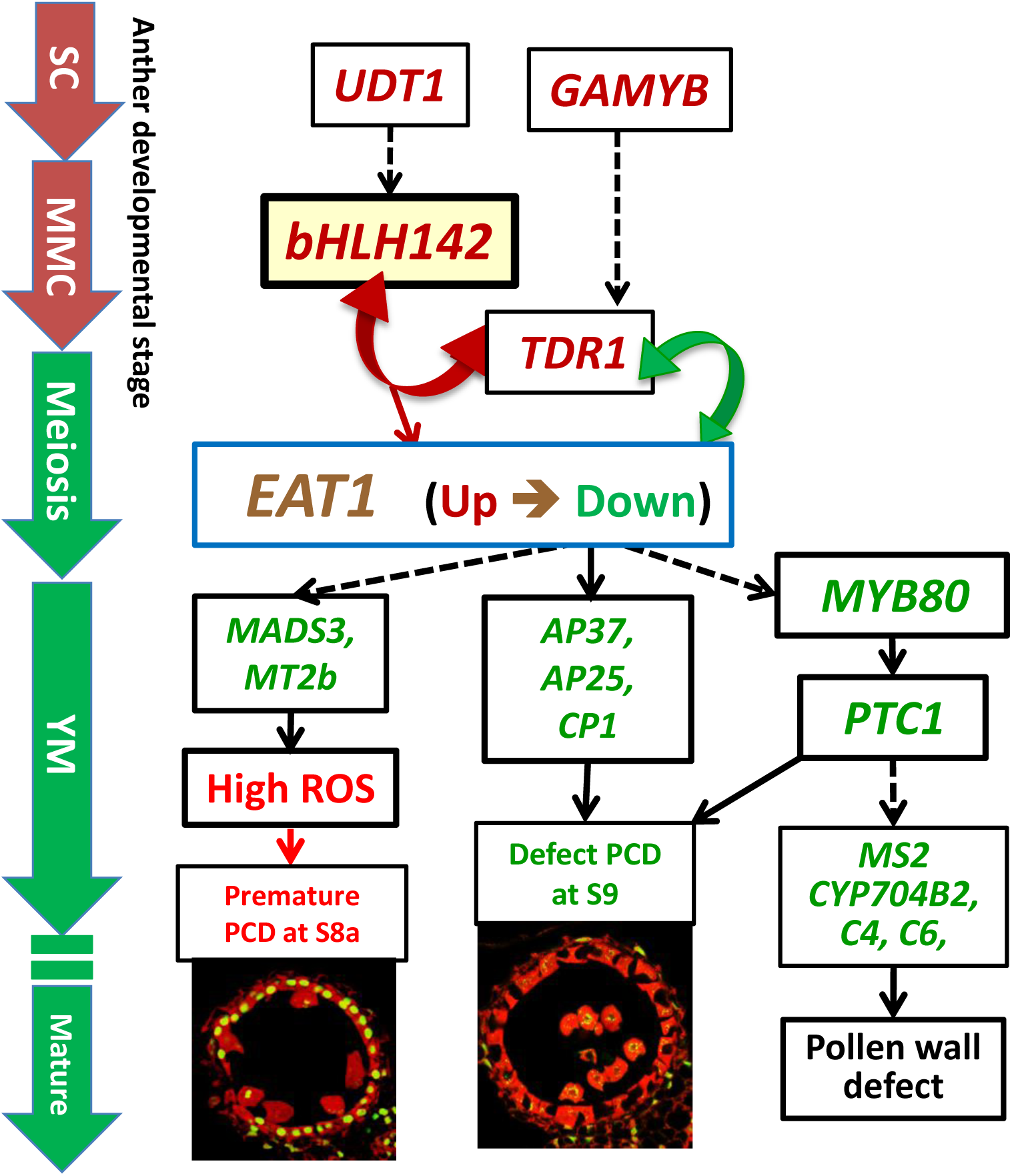
Proposed mechanistic model of male sterility in rice caused by overexpressing *bHLH142*. Overexpression of *bHLH142* upregulates *UDT1, GAMYB, TDR1*, and *EAT1* at an early stage of anther development. However, *EAT1* is downregulated at the young microspore stage (YM, S9), which in turn reduces the expression of the downstream genes involved in ROS scavenging *(MADS3, MT2b)*, leading to premature onset of tapetal PCD at meiosis-I (S8a). Moreoer, OE142 downregulated PCD *(AP37, AP25, CP1)*, and sporopollenin biosynthesis *(MYB80, PTC1, MS2, CYP704B2, C4, C6).* Thus, defected tapetal PCD at YM stage (S9) and defected pollen wall development together leads to male sterility in the overexpression line. Genes marked in red or green denote upregulation or downregulation, respectively. Solid arrow lines indicate direct regulation, while dotted arrow lines indicate indirect regulation. Double arrows represent protein-protein interaction. SC, sporogenous cell; MMC, microspore mother cell; YM, young microspore stage (S9).

Western blot analysis further revealed that bHLH142 protein was specifically accumulated in the OE142 anthers (**Fig. 3F**). EAT1 protein was not detectable at S8a but accumulated at a high level at S9 in the Wt (S9). However, OE142 anthers precociously expressed EAT1 protein at S8a but its expression was reduced to negligible level at S9 (**Fig. 9C**). These data suggest that overexpression of *bHLH142* prematurely upregulate *EAT1* transcription (**Fig. 5F, 9A**) as well as its protein level (**Fig. 9C**) in the anthers of OE142. Thus, the decreased transcript abundance and protein amount of EAT1 at S9 might interrupt the normal anther development in the OE142 transgenic lines.

In our previous study, we demonstrated that bHLH142 interacts with TDR1 to co-modulate *EAT1* transcriptional activity (Ko *et al.*, 2014). Overexpression of *bHLH142* increases bHLH142 protein level, which may in turn enhance bHLH142/TDR1 protein-protein interaction to increase *EAT1* expression at the early stage of anther development. However, downregulation of *TDR1* expression at S8 and onward **(Fig. 5E)** might decrease TDR1 protein translation and then hamper TDR1/bHLH12 protein-protein interaction, therefore significantly reduce *EAT1* expression at later stages even though high bHLH142 protein is present in the anthers of OE142. In additions, a low EAT1 protein level at S9 **(Fig. 9C)** might further reduce TDR1/EAT1 interaction and subsequently influence the regulatory cascade of downstream target genes and result in defected pollen development in OE142 anthers.

## DISCUSSION

### Overexpression of bHLH142 causes male sterility by triggering premature PCD

In an effort to provide greater insight into the functionality of bHLH142 in rice pollen development, we generated transgenic lines constitutively overexpressing *bHLH142.* To our surprise, overexpression of *bHLH142* also leads to male sterility in rice, similar to the knockout mutant reported previously (Ko et al., 2014). Except for the defect in pollen development, OE142 transgenic lines maintain wild-type like vegetative growth (**Fig. 1; Supplementary Fig. S1, S2**).

Our in-depth molecular characterization suggests that overexpression of *bHLH142* significantly alters *in vivo* homeostasis of the known key pollen development-related regulatory TFs in the OE142 anthers (**Fig. 5**). For example, *UDT1, GAMYB, MYB35, TDR1*, and *EAT1* were upregulated in OE142 at the early stages of anther development (Fig. 5). Presumably, this triggers a premature onset of tapetal PCD in OE142 anthers before the maturation of pollen grains, as shown in the TUNEL assay (**Fig. 2**). The reduced expression of *EAT1* at the young microspore stage (S9) and onward in OE142 anthers (**Fig. 5F**) further impairs the normal development of pollen grains because of reduced expression of the downstream genes in the PCD pathway, such as *AP37, AP25*, and *CP1* (**Fig. 6**) and pollen wall biosynthesis, such as *MYB80, PTC1, MS2, Cyp704B2, C4*, and *C6* (**Fig. 8**). Obviously, interference with the timely expression of these pollen development associated genes leads to male sterility in OE142.

### Tightly regulated bHLH TFs are essential for pollen development

Our study revealed that overexpression of *bHLH142* causes significant changes in the expression of known regulatory genes associated with tapetal PCD, ROS metabolism, and pollen wall development (**Fig. 6–8**), thus leading to male sterility in OE142 transgenic plants. This may result from the interference in its protein interaction with TDR1 in activational transcription of *EAT1 (Ko et al., 2014).* This study advances our knowledge of the molecular mechanism underlying the bHLH142 and EAT1 transcriptional circuits controlling pollen development in rice and possibly in other plants as well. Timely expression and maintenance of proper expression levels of these bHLH TFs must be tightly regulated developmentally for normal pollen maturation. The hierarchy of several known regulatory network genes associated with pollen development is therefore clarified in this study.

Based on this and previous studies, we propose a mechanistic model of genic male sterility in rice as caused by overexpressing *bHLH142* (**Fig. 10**). According to the model, overexpressing *bHLH142* causes upregulation of *UDT1, GAMYB, TDR1*, and *EAT1* at the early stage of anther development. This consequently leads to premature onset of tapetal PCD. However, *EAT1* is downregulated at the young microspore stage (YM, S9) in OE142 anthers, which in turn further reduces the expression of the downstream functional genes involved in PCD (*AP37, AP25*, and *CP1*), ROS scavenging (*MADS3* and *MT2b*), and pollen wall biosynthesis (*MYB80, PTC1, Cyp704B2, MS2*, and *C4*) and impairs normal pollen grain maturation. Thus, increased ROS accumulation, defect in timely tapetal PCD at the YM stage, and defect in pollen wall development, eventually lead to male sterility in the OE142 plants. The alterations in homeostasis of key TFs in pollen development or protein-protein interaction between bHLH142/TDR1 or TDR1/EAT1 may account for the decreased expression of downstream pollen development marker genes regulated by EAT1 (Ko *et al.*, 2014).

### Potential of establishing a male sterility line by overexpressing key TFs

Our finding that overexpression of *bHLH142* (*TIP2*) causes male sterility by triggering premature PCD in rice is similar to previous results obtained by overexpressing several pollen development related transcription factors in other species. A total of 148 out of 196 *Arabidopsis* transformants overexpressing *AMS* (rice homolog of *TDR1)* produced sterile pollen, mimicking the *ams* mutant phenotype (Sorensen et al., 2003). It is claimed that the resulting male sterility might be due to co-suppression of *AMS*, but our results tend to suggest that altered homeostasis of the related TFs may be the major cause. Moreover, overexpressing *MS1* as driven by the CaMV35S promoter also caused stunted plants with sterile pollen in *Arabidopsis* (Yang *et al.*, 2007).

Recently, the ortholog of *MS1* in barley *(HvMS1)* was cloned and its expression was altered to be either overexpressed or suppressed (Fernandez Gomez and Wilson, 2014). Both RNAi and overexpression of *HvMS1* full-length cDNA under the control of the maize ubiquitin promoter caused male sterile phenotype in the transgenic barley plants. Also, knockout of *AtCEP1*, which encodes a papain-like cysteine protease involved in tapetal PCD, delayed tapetal PCD, while its overexpression caused premature tapetal PCD (Zhang *et al.*, 2014). Thus, we hypothesize that overexpression of other key TFs in the pollen development regulatory network, such as *GAMYB, UDT1, TDR1*, or *EAT1 (DTD1, bHLH141)* may also cause male sterile phenotype in rice due to alteration in the homeostasis of the regulatory cascades in pollen development.

### Advantages of using OE142 in hybrid seed production

In this study, overexpression of *bHLH142*, an anther-specific TF gene, by a strong constitutive promoter led to its ubiquitous transcription in leaves, hulls, as well as in the anther of OE142, as expected (**Fig. 3**). However, bHLH142 protein expression was not constitutively expressed. Rather, its expression was maintained in a tissue specific manner in the anthers **(Fig. 3F** and **4)**. The ubiquitin promoter is expected to drive ubiquitous gene expression. However, protein expression level is determined by the rate of transcription and by post-transcriptional processes that lead to changes in mRNA transport, stability, and translational efficiency. The nuclear localization sequence (NLS) mediates the transport of nuclear proteins into the nucleus. bHLH142 contains two NLS and targeted protein is localized in the nucleus (Ko et al., 2014). The basis for the specific expression of bHLH142 in anthers, as driven by the ubiquitous promoter, requires further study. Some unknown posttranscriptional factor(s) specifically in the anthers may be required for its translation. In fact, overexpressing target genes in an anther-specific manner is desirable from the perspective of GMO food biosafety because anther-specific expression will avoid any unintended expression in other tissues, especially in the edible part of the seed **(Fig. 4)**. This may be more acceptable to consumers.

### Future prospects

Cytoplasmic male sterility (CMS) is widely used in F1 seed production in rice. Wild abortive (WA) CMS gene encodes a tapetal mitochondrial protein, WA352, which interacts with the nucleus encoded mitochondrial protein COX11 to inhibit ROS-scavenging activity, thus triggering premature tapetal PCD and pollen abortion (Luo *et al.*, 2013). Whereas CMS guarantees a high degree of purity of hybrid seeds it increases production cost. Therefore, there is a great demand for the generation of new male sterility lines for production of hybrid crops, such as rice, maize and wheat, etc. Here, we showed that overexpression of *bHLH142* may provide a novel and simple way to generate genic male sterility lines in rice. Normally, genetic engineering using the overexpression approach is preferred to RNAi by the biotech industry. However, one major concern with this technology is the maintenance of the genic male sterility line as it fails to produce viable seeds as stock. A strategy to generate inducible male sterility and restorer hybrid seed production has been demonstrated in *Arabidopsis* (Li *et al.*, 2007). In this system, a complete male sterility transgenic line was generated using AtMYB103EAR chimeric repressor construct under the control of the native AtMYB103 promoter, whereas the restorer containing the AtMYB103 gene under the control of a stronger anther-specific promoter (p39) was introduced into the pollen parent (Li *et al.*, 2007). Fertility is restored when the restorer line is crossed with the male sterile plant. Another possible solution to alleviate this problem is to employ an inducible promoter to drive its expression. GM male sterility plants generated by overexpression of a MYC5-SRDX chimeric repressor under the control of an inducible promoter have been demonstrated in *Arabidopsis* (Figueroa and Browse, 2015). Other promoters inducible by chemicals, temperature or light can also be considered and tested.

## Accession numbers

Sequence data from this article can be found in the GenBank/EMBL database under the following accession numbers: bHLH142 (Os01g0293100), protein (NP_001042795.1). Additional loci are presented in **Supplementary Table S1**.

## Acknowledgments

We thank the Tos17, Postech, and TRIM Mutant Libraries for providing rice mutant seeds. We appreciate the technical support of the Biotechnology Center in Southern Taiwan, Academia Sinica, GMO greenhouse core facility. This work was supported in part by the Biotechnology Center in Southern Taiwan, Academia Sinica, and a Ministry of Science and Technology of Taiwan grant to Dr. Swee-Suak Ko’s project (MOST104-2313- B-001 -003). We thank Ms. Miranda Loney for English editing.

## Supplementary data

**Supplementary Table S1.** Primers used in this study.

**Supplementary Fig. S1.** *bHLH142* overexpressing transgenic lines.

(A) Panicles of wild-type (Wt) and different OE142 T_0_ lines at the mature stage. (B) Spikelets of Wt (left) and several OE142 T_0_ lines at one day before anthesis. (C) Grains of Wt (filled) and several OE142 T_0_ lines at the harvest stage. (D) De-hulled seeds of Wt (filled) and several OE142 T_0_ lines showing inviable seeds at the harvest stage. (E) Construct map and primer design for genomic PCR to confirm T-DNA insertion. (F) Genomic PCR confirmed T-DNA insertion in different OE142 lines.

**Supplementary Fig. S2.** Overexpression of *bHLH142* (OE142) resulted in defect in anther development in rice.

(A) Staining of anther and pollen grains by 2% I_2_/KI solution in the wild-type (Wt) and OE142 line #96 at 1 day before anthesis (DBA). (B) Weak staining of Sudan Black in the transverse anther section of OE142 (right panel) compared to the Wt (left panel) at 1 DBA. (C) Transverse anatomical comparison of anther of the wild-type (Wt) and OE142 at 1 DBA using DIC. Arrows show the fertile pollen (B, C) and the thickening of endothelial cell layers, prior to anther dehiscence (C). Arrowheads show degenerated pollens (C, right panel). Scale bars: 100 um (A), 20 um (B, C).

**Supplementary Fig. S3.** Transverse sections showing defect of anther development in OE142 line.

Arrows indicate the thin epidermal layer in OE142 anther. Arrowheads show the degenerated pollens. Scale bars: 20 μm.

**Supplementary Fig. S4.** Differential interference contrast (DIC) images of anther cross sections corresponding to TUNEL assay.

Upper panel: wild type, lower panel: OE142. Scale bars: 50 μm.

**Supplementary Fig. S5.** Mutagenesis analysis indicated gene hierarchy of *MYB80* and *PTC1.* Wild type for *tdr1* is Dongjin, *eat1* is in Hitomebore background. Abbreviations are as described in the legend of Fig. 5.

**Supplementary Fig. S6.** RNA *ISH* to *EAT1-* sense probe to anthers at various developmental stages of the Wt and OE142.

Scale bars: 20 μm.

